# Sensory Appendage Protein triggers alarm to pyrethroid in Indian malarial vector Anopheles culicifacies

**DOI:** 10.1101/2025.01.21.634214

**Authors:** Vaishali Saini, Pooja Rohilla, Vartika Srivastava, Gitanjali Tandon, Pooja Yadav, Nirmala Sankhala, Tanvi Singh, Gunjan Sharma, Suchi Tyagi, Tanwee Das De, Rajnikant Dixit

**Affiliations:** Laboratory of Host-Parasite Interaction Studies, Department of Vector Genomics, ICMR-National Institute of Malaria Research, Dwarka, New Delhi, 110077, India; Academy of Scientific and Innovative Research (AcSIR), Ghaziabad, Uttar Pradesh-201 002; Department of Medical Entomology and Zoology, ICMR-National Institute of Virology, Microbial Containment Complex, Pune

**Keywords:** Anopheles culicifacies, Sensory Appendage proteins, Mosquito host-seeking, Insecticide, Behavioural responses

## Abstract

Sensory appendage proteins (SAPs) serve as molecular arms of odorant-binding protein family members and mediate chemical communication from the external to the internal environment of the insect body. We recently reported that SAP members might have an important role in blood-feeding and/or insecticide resistance-associated behavioral physiologies, but the mechanism remains unclear. Here we show how *SAP* contributes (i) to the host attraction during navigation for blood-feeding, and (ii) triggers an alert *via* integrated actions of eyes and peripheral sensory organs to circumvent the toxic effect of insecticide in the susceptible strain of *An. culicifacies*. Among the six identified SAP members, *AcSAP1* and *AcSAP2* are abundantly expressed in the olfactory and legs, and their expression is significantly modulated at night in naïve adult female mosquitoes. We noticed that *AcSAP1* silencing significantly impairs the host-attraction properties of the mosquitoes. Its strong binding affinity to the synthetic pyrethroid prompted us to evaluate the *AcSAP1* response to insecticide exposure. Confocal microscopy and molecular profiling data reveal a heightened modulation in the expression of *AcSAP1* in eyes, and other peripheral sensory organs including the olfactory (****p<0.00001), legs (**p < 0.004), wings (***p < 0.0004). These findings collectively suggest a crucial role of *AcSAP1* in facilitating mosquito navigation, and/or insecticide avoidance. Unraveling the basis of SAP interactions with peripheral sensory systems could pave the way for the development of novel vector control tools.

## Background

To feed, adapt, and survive under different environmental conditions, mosquitoes have developed an efficient and sophisticated sensory system. They rely on their sense of smell to navigate their surroundings and find appropriate food sources, identify mates, and ovipositing sites. The primary source of nourishment and energy for both male and female mosquitoes is nectar sugar (1). However, female mosquitoes consume blood to reproduce and maintain the gonotrophic cycle. During the blood foraging states, a mosquito’s host-seeking behavior is dominantly guided by the controlled action of the neuro-olfactory system (2). Comprehending the molecular basis of mosquito olfactory sensory modalities may open new avenues for the development of malaria control strategies.

The olfactory system encodes a diverse array of olfactory proteins and receptors which synergistically coordinate to facilitate navigation (3). Two major categories of small soluble proteins i.e. Odorant-binding proteins (OBPs), and chemosensory proteins (CSPs), are believed to play a dynamic role in the chemoreception process which is largely mediated by odorant receptors (ORs), ionotropic receptors (IRs), and sensory neuron membrane proteins (SNMPs). Accumulating evidence highlights OBPs, which are small soluble, freely circulate in high concentration within the chemo-sensilla lymph, and carry a unique ability to bind and transport hydrophobic stimuli (odorants, pheromones, etc.) to chemoreceptors present on dendritic neurons (4). Thus, OBPs serve as an intermediary between surrounding odorants and their receptors, modulating various mosquito behavioral properties such as host-seeking, and blood feeding. Like OBPs, chemosensory proteins (CSPs), also known as Sensory Appendages Proteins (SAPs), are small soluble hydrophobic proteins containing OS-D domains, however, their defined role remains poorly investigated in mosquitoes.

Limited studies highlight that the *Anopheles* mosquito genome encodes at least 6 transcripts (CSP1-CSP6), which are predominantly expressed in the olfactory system (5). Our previous RNA-Seq analysis also identified similar proteins, of which two members of CSPs i.e. SAP-1 (ACUA018714) and SAP-2 (ACUA013409) were abundantly expressed in the legs and olfactory tissues. Their rhythmic fluctuation during late night suggests their role in the onset of host-seeking behavioral activities of the mosquito *An. culicifacies* (3). Later, a study not only confirmed the expression of SAP in the legs but also demonstrated their involvement in insecticide resistance tolerance in mosquito *An. gambiae* (6). However, the underlying mechanism remains undefined.

*An. culicifacies* transmits approximately 65% malaria in rural India (7), but significant knowledge gaps linked to the resistance level to the most effective pyrethroid-based LLINs, or IRS tools, and their effect on behavioral physiology which poses a challenge in controlling this vector population. To evaluate the possible role of the newly identified SAP in the host-seeking activity and insecticide resistance we conducted detailed transcriptional profiling and functional studies in the mosquito *An. culicifacies*. Here, we demonstrate that SAPs not only influence host-seeking associated behavioral properties but also trigger rapid hyper-sensitization of the body’s peripheral sensillum organs in response to pyrethroid exposure. Our data provides substantial evidence for future exploration of the SAP’s role as a warning against toxicants, a knowledge that could be translated into strategies to combat widespread pyrethroid resistance in the wild mosquito population.

## MATERIALS AND METHODS

### Mosquito Rearing and Maintenance

*An. culicifacies* (sibling species A) was reared and maintained in the central insectary of the National Institute of Malaria Research under standard rearing conditions of 28 ± 2°C, relative humidity 60–80%, and 12:12 hr light/dark cycle. The aquatic stages were reared in enamel trays using water supplemented with a mixture of dog food and fish food. After emergence, adults were kept in mosquito cages and fed on cotton swabs dipped in 10% sugar solution. All protocols for mosquito rearing and cyclic colony maintenance were duly approved by the institute’s ethical committee [NIMR/IAEC/2017-1/07] (8).

### Tissue collection and RNA extraction

Various tissues such as olfactory, brain, legs, wings, and eyes were dissected from the ice-anesthetized adult female *An. culicifacies* mosquitoes’ and collected in Trizol Reagent (RNAiso Plus, Takara Bio, Kusatsu, Japan). Developmental stages of *An. culicifacies viz*. egg, larvae (stages I − IV), and pupae were also collected in Trizol after excess water was removed using filter paper. Total RNA was isolated by the standard Trizol method as described previously, (9) and quantified by using a Nanodrop 2000 (ND-2000c) spectrophotometer (Thermo Scientific, USA).

### cDNA preparation and gene expression analysis

Approximately 1μg of total RNA was used to synthesize the first strand of cDNA using Primescript^TM^ 1^st^ strand cDNA Synthesis Kit (Takara Bio, Kusatsu, Japan) following manufacturer’s protocol. Routine differential gene expression analysis was performed by the RT-PCR. Relative gene expression analysis was analysed using SYBR green qPCR (Takara, Bio Inc, Kusatsu, Japan) master mix using a Bio-Rad real-time machine (CFX96^TM^ Real-Time system, USA). The four-step PCR cycle included an initial denaturation at 95°C for 5min, followed by 40 cycles of 10s at 95°C, 15s at 52°C, and 22s at 72°C. Fluorescence readings were recorded at 72°C after each cycle. In the final steps, PCR was conducted at 95°C for 15s, followed by 55°C for 15s, and again 95°C for 15s, before generating a melting curve (10). The reproducibility of the result was ensured by repeating the experiments with three independent biological replicates. Throughout the experiment, the *actin* gene was used as an internal control, and relative quantification data were analysed by 2^−ΔΔCt^ method (11). Relative expression graphs were plotted using GraphPad Prism 10 and the significance was determined by multiple comparisons using one-way ANOVA.

### dsRNA-mediated gene silencing

To test the effect of *AcSAP1*-knockdown, an *in vitro* transcription reaction was performed to prepare dsRNA by using Transcript Aid T7 high-yield transcription kit (Cat# K044, Ambion, USA). Approximately 69nl of purified dsRNA at a concentration of ∼3μg/μl for the target gene was injected into cold anesthetized 1–2-day(s)-old *An. culicifacies* female mosquitoes using a Nano-injector (Drummond Scientific, CA, USA). For the control group, age-matched mosquitoes were injected with bacterial origin *LacZ* dsRNA. Three days post dsRNA injection, both control and experimental groups of mosquitoes were used for phenotypic assays. Simultaneously, to examine the silencing efficiency, olfactory and leg tissues were collected in Trizol from 20 mosquitoes each from the control and SAP-silenced mosquitoes for RNA extraction and cDNA preparation. The efficiency of silencing was evaluated using quantitative real-time PCR as described above.

### Host-seeking behavioral assay

To assess the behavioural and phenotypic effect a host-proximity assay was carried out to compare the host-seeking efficiency between control and *AcSAP* silenced mosquito groups. Human host proximity assay was performed by minor modification of previously described methods (12). The two-port mosquito behavioural assay chamber was designed with dimensions of 30 cm in height, 40 cm in width, and 30 cm in length. It was fabricated using flexible glass material. The chamber includes a central splitting door, creating two equal-sized compartments, each with dimensions of 30 cm in height, 20 cm in width, and 30 cm in length. One compartment is designated for the control mosquitoes’ group, while the other is for knock-down mosquitoes. Each chamber is built with an arm (L×D: 27×12cm) for placing human hands.

Before the commencement of the behavioural experiment, 25 non-blood-fed mosquitoes from both the control and the SAP-silenced group were released into their respective chambers and allowed to acclimatize for 5-10 minutes. After acclimatization, the hands of the volunteer were placed into the arms of the behavioural assay chamber. The usage of soap or any cosmetic products was restricted for the volunteers participating in this assay. A gauze barrier was incorporated into the arm (at 15cm) and the distance between the hand and the barrier was 0.5cm to avoid direct landing and biting by mosquitoes on the hand. The protocol was duly approved by the Institute Biosafety Committee (IBSC/NIMR/2022/06).

Considering the nocturnal behavioural activity, the assay was conducted at ZT16-17, and data was recorded manually for 8-10 minutes. The number of mosquitoes that responded and attracted to the hands was counted manually. At an interval of every 30sec-1min, we counted the number of mosquitoes in real-time, that were attracted to the volunteer’s hands, landed on the gauze, and attempted to bite. The difference in host-seeking efficiency was calculated as the percentage of attracted mosquitoes by quantifying the ratio of landed and total number of mosquitoes used in the respective assay. The experiment was repeated three times (three cohorts of mosquitoes).

### In-silico prediction of SAP binding affinity with deltamethrin

The protein sequences of SAP1, SAP2, and SAP3 were subjected to physiochemical analysis with the help of ProtParam tool (https://web.expasy.org/protparam/) (13) at EXPASY while the Cellular localization of SAP proteins was predicted by using web servers CELLO v.2.5 (http://cello.life.nctu.edu.tw/) (14) and WoLF PSORT (http://wolfpsort.seq.cbrc.jp/) (15) To carry out protein-ligand docking, firstly three-dimensional structures of all three SAP proteins were downloaded from the AlphaFold Protein Structure Database (https://alphafold.ebi.ac.uk/) (16). To check the quality of the 3D structures of the SAP proteins, PROCHECK was done using SAVES server (https://saves.mbi.ucla.edu/) (17). Using these structures, SAP proteins and Deltamethrin docking was done with the help of Autodock Vina (https://vina.scripps.edu/) (18). Later the obtained docked complex was further analysed using PyMol (https://pymol.org/) (19) and BIOVIA Discovery Studio v24.1.0.23298 (https://www.3ds.com/products/biovia/discovery-studio) (20).

### Insecticide Exposure and SAP Response

Healthy, naïve 3–4-day-old adult females of *An. culicifacies* were exposed to World Health Organisation (WHO) recommended insecticide pre-impregnated papers containing deltamethrin (0.05%), obtained from University Sains Malaysia, Malaysia. Control tests were conducted using pre-impregnated paper with silicone oil (as a deltamethrin control) alongside insecticide. For each test, 25 mosquitoes were exposed for 1 hour, and cumulative knock-down was recorded at 10-minute intervals. The mosquitoes were then transferred to a holding tube and fed a 5% sucrose solution. The survival test was conducted in the laboratory at the prescribed temperature (27±2°C) and humidity conditions (75 ± 10%). After 1 hour of exposure, both dead and alive mosquitoes were counted, and mortality in the test and control replicates was determined to generate the survival curve (21). To examine molecular responses to insecticide exposure, desired tissues were collected from the surviving mosquitoes, and SAP expression was evaluated by using a relative quantification assay as described above.

### Immunofluorescence and co-localization assays

To validate the molecular expression data we first generated polyclonal antibodies against *in-house* produced recombinant protein (see Supplementary Text; and **Fig: S4**) and performed immunofluorescence assay for cellular localization. 3–4-day-old adult female mosquitoes *An. culicifacies* were fixed in 4% formaldehyde in PBS at room temperature for 20 minutes. For immunolabeling, the mosquito body and/or targeted tissue was permeabilized for 1 h in 0.01% Triton X-100, saturated in 1% BSA for 1 h, treated with a solution of the first antiserum overnight, and then stained with a solution of the fluorescein-conjugated anti-rabbit IgG antiserum 1:80 for 1 h. All solutions were prepared in 1X PBS and all treatments were performed at room temperature. Three washes of 5 min each with 1X PBS were included between all the treatments and at the end of the procedure (22). The samples thus prepared were mounted on microscopic slides and observed under a confocal microscope (Olympus FLUOVIEW FV1000).

## Results

### Age and circadian rhythm modulate SAP expression

From our recent RNAseq study, we identified at least seven putative chemosensory proteins including three Sensory Appendages Proteins (SAP1, SAP2, and SAP3; Supplementary Table-1), which are low molecular weight (∼12-14k Da) small peptides expressing in the olfactory tissue of the naïve 3–4-day old adult female mosquito *Anopheles culicifacies* (3). But their role in mosquito behavioral physiology remains unexplored, and hence, first, we carried out a detailed spatial/temporal expression analysis under the distinct physiological status. Initially, we noticed that all putative CSPs/SAPs were duly expressed during the aquatic developmental stages of the mosquitoes (Supplementary Fig:S1, S2). Comparative to other SAPs/CSPs, an observation of higher expression of *AcSAP1* in the sensory tissues (Fig. 1a), endorsed us to further decode its role in host-seeking associated behavioral modification properties in the adult female mosquito. Age-dependent expression data revealed a gradual increase in the *AcSAP1* till 3-4 days, which significantly downregulated in the olfactory system of 9-10 days-old female mosquitoes (Fig1b). However, we also noticed late-night elevated expression in 3-4 days adult females (Fig1c.), corroborating with the previous observations (3).

**Figure 1:**
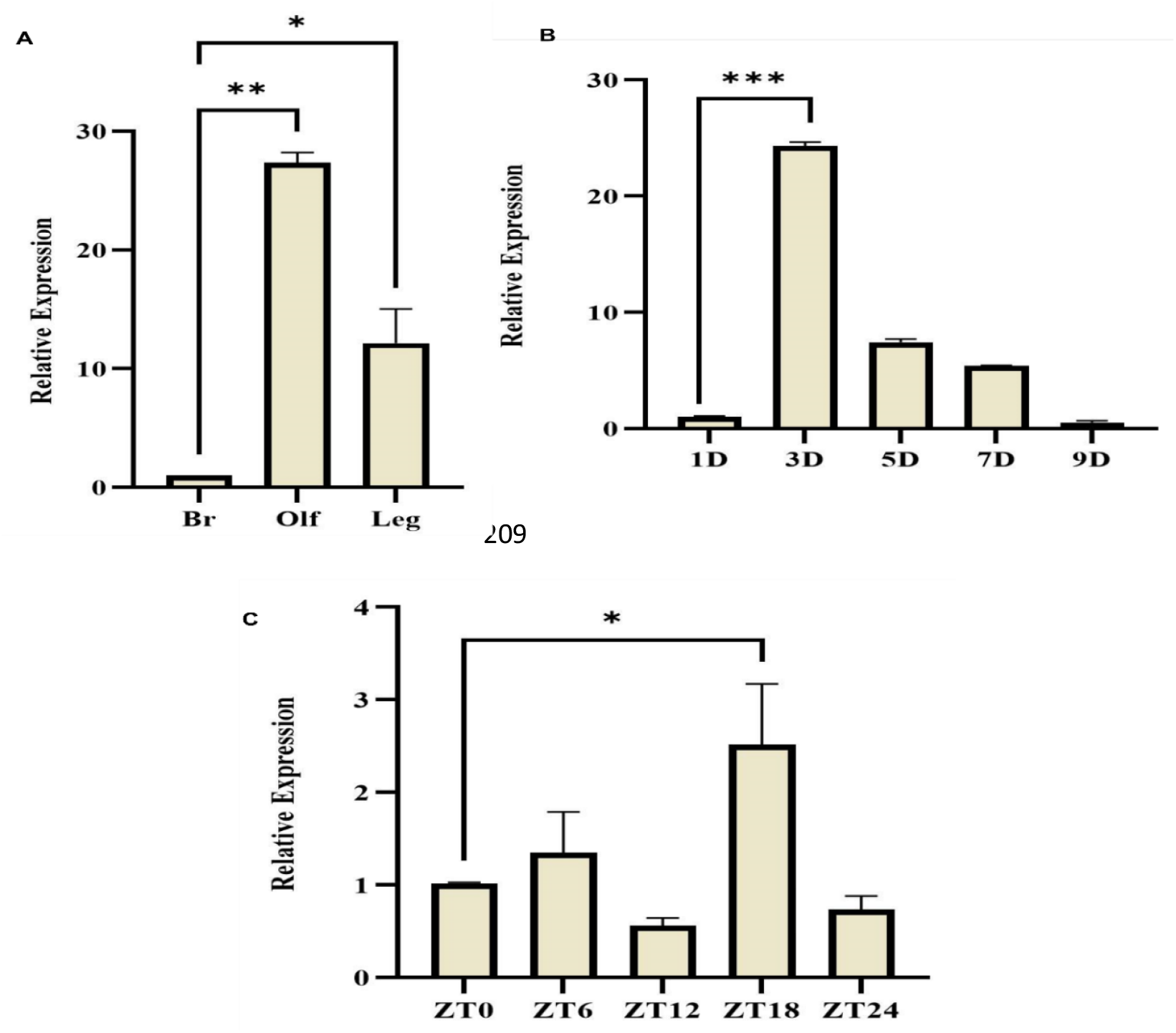
Spatial and temporal expression profiling of *AcSAP1*: **(A)** *AcSAP1* dominantly expressed in the olfactory (Olf) and leg (Leg) of the adult female mosquitoes: Tissues were collected from 3– 4-day-old females from mixed cohort and compared with female brain as a control *(n = 25, N=3);* **(B)** Age-dependent **(D= days)** transcriptional profiling of *AcSAP1* in the non-blood-fed adult female mosquito’s olfactory system; **(C)** Zeitgeber time (ZT) scale dependent rhythmic expression of *AcSAP1* protein in the adult female’s olfactory system (OLF) where ZT0 indicate the end of dawn transition, ZT11 is defined as the start of the dusk transition and ZT12 is defined as the time of lights off. All the three independent biological replicates statistical analysis was performed by one-way ANOVA where asterisks represented *p < 0.01; **p < 0.001; and ***p < 0.0001, (n = represents the number of mosquitoes pooled for sample collection; N = number of replicates).

**Figure 2:**
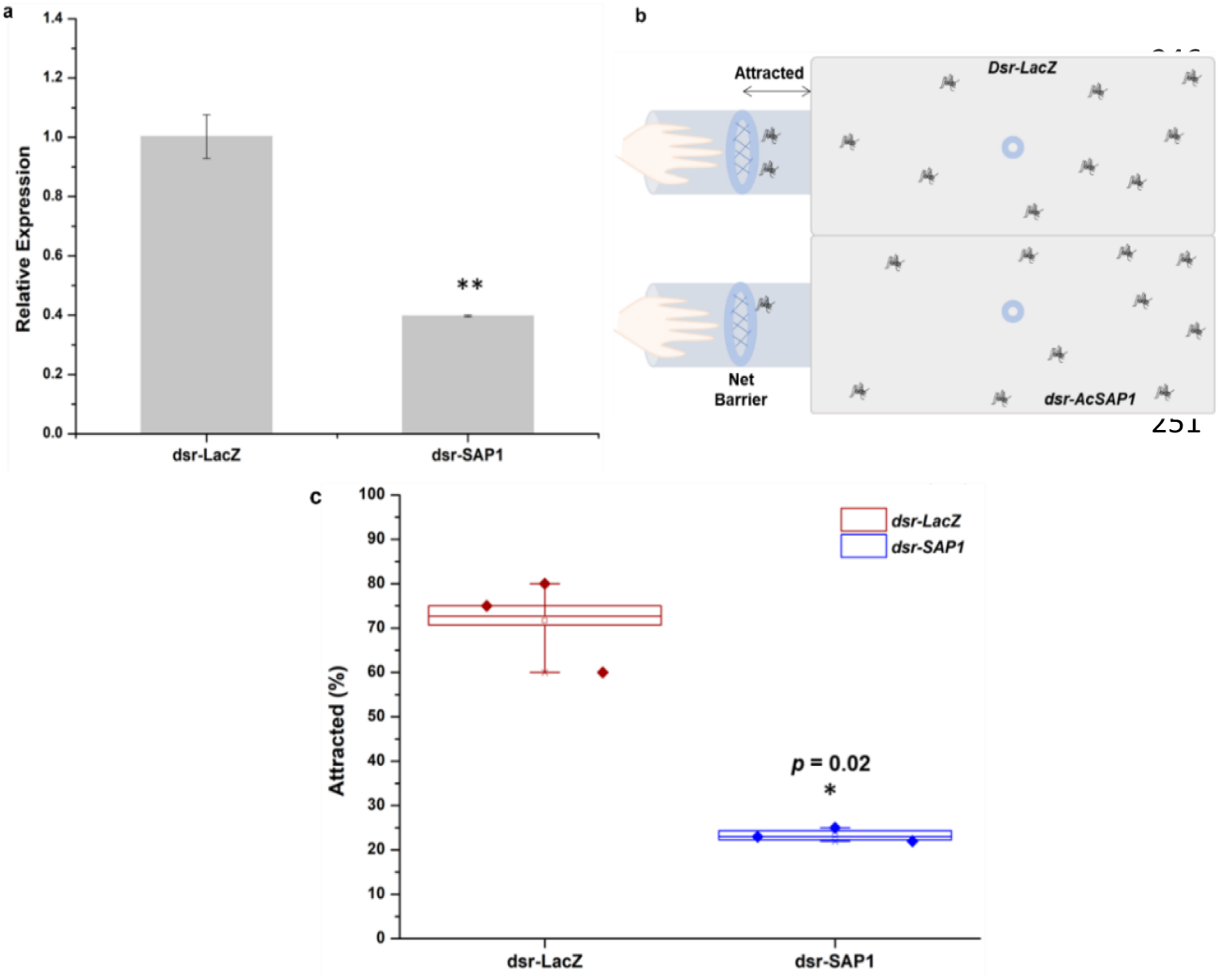
*AcSAP1* silencing interferes with host attraction properties: **(a)** Fold change analysis of silencing validation of *AcSAP1* expression in the olfactory system; **(b)** Schematic representation of host-seeking efficiency assay; **(c)** comparative analysis of mosquitoes’ attraction to human host in dsr-*LacZ* treated and dsr-*AcSAP1* injected mosquitoes. *AcSAP1* knock-down mosquitoes showed significantly lower attraction compared to their control counterpart (p = 0.02, χ 2 test).

### AcSAP1 influences host attraction properties

An enriched expression of *AcSAP1* in the olfactory tissues and legs of adult female *An. culicifacies* mosquitoes led us to hypothesize that *AcSAP1* plays a critical role in the pre-blood-meal associated behavioral property of host attraction. To test this, adult female mosquitoes, both control and *AcSAP1* silenced groups, were given a 10-minute window to seek out a host for feeding. We compared the number of mosquitoes attracted during this period.

We noticed that *AcSAP1-silenced* mosquitoes exhibited a delayed activation of approximately 2.5 minutes compared to the control group. By the end of the recording period, at least 70 ± 10% of the control group mosquitoes were attracted to the host, while the silenced group showed a significant reduction, with 23 ± 2% fewer mosquitoes being attracted. Overall, our data highlights a delayed attraction of 65 ± 6% in silenced mosquitoes compared to the control, suggesting that *AcSAP1* is crucial for regulating the host attraction behaviour in *An. culicifacies*.

### Insecticide exposure modulates AcSAP1 expression

The unavailability of the insecticide-resistant strain, inducted us to evaluate the possible roles of *AcSAP1* in the insecticide-mediated behavioral modification in the laboratory-reared susceptible strain of mosquito *An. culicifacies*. Initially, we performed a detailed *in-silico* prediction and molecular docking assays of all three SAPs with pyrethroid deltamethrin. Interestingly, we noticed all three SAPs had similar molecular features though a variation was readily observable in their physiochemical properties (Supplementary Table 2). Analysis of the predicted 3D structure and Ramachandran plot revealed that all three SAP proteins have most of the atoms lying in the core regions, limiting to very few or none in the disallowed region (Fig A; Supplementary **Fig. S3**). However, we also noticed all three SAPs meets the quality criteria for the *in-silico* binding however docking analysis further inferred that *AcSAP1* possessed the greatest binding affinity i.e. -5.3 kcal/mol as compared to SAP2 and SAP3 (Fig. 3B; **Fig S3**). Next, we performed WHO bottle assay to further test and validate the *in silico* predicted properties of *AcSAP1*. As expected, we observed a maximum survival till 30 minutes of the susceptible strains, when exposed to Deltamethrin (Fig.3 C-D). Finally, a multi-fold enrichment of SAP expression in all the tested peripherally active tissues viz. Olf, Legs, and Wings confirmed that *AcSAP1* has a higher ability to bind to Deltamethrin (Fig. 3. E-G).

**Figure 3:**
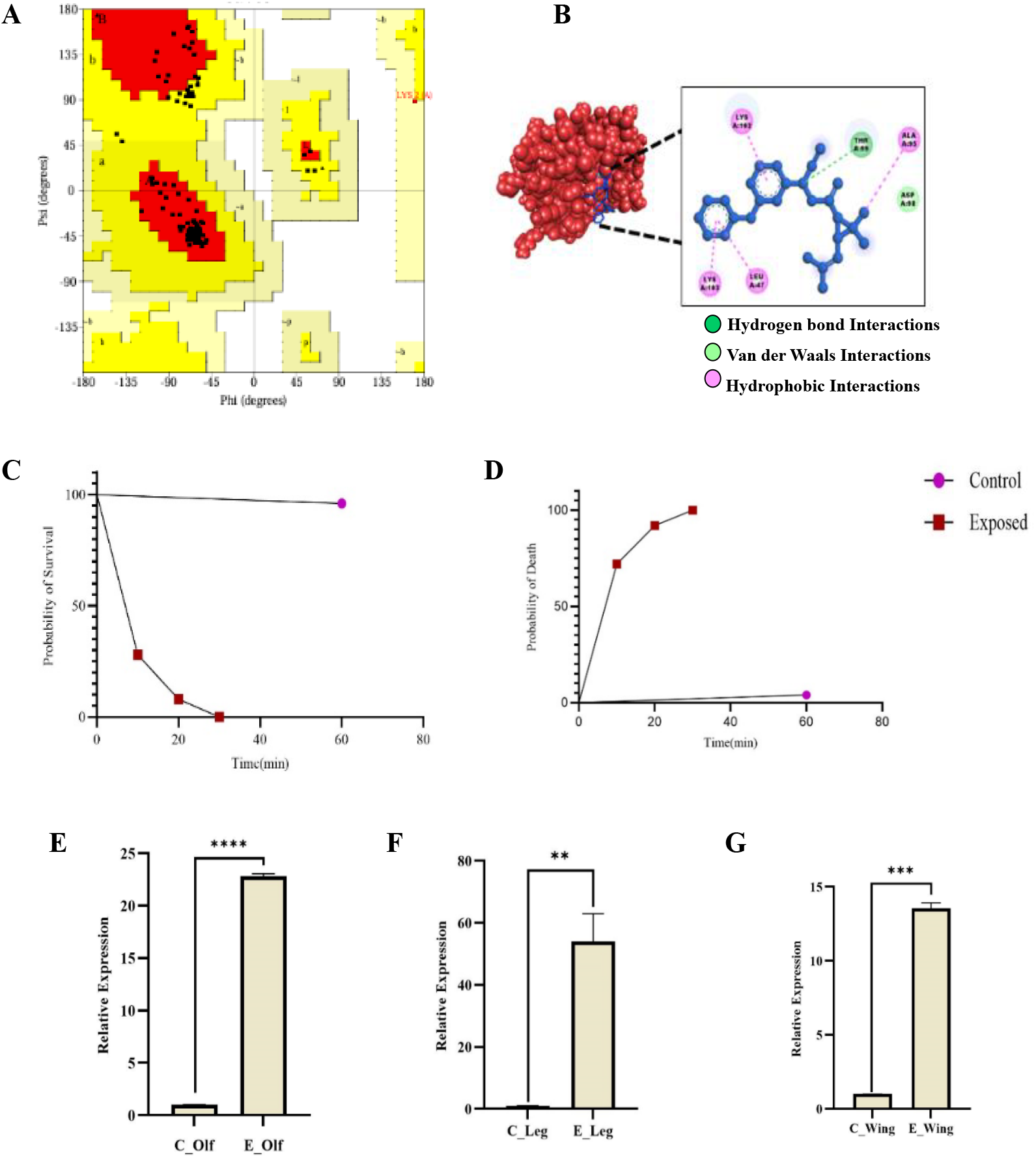
*In-silico* structural analysis of SAP-DM binding: **(A)** Ramachandran Plots showing SAP proteins favors DM binding as most of the atoms lie in the core regions, (also see Supplementary Fig.S3). **(B)** Docked complexes of SAP proteins with Deltamethrin showing SAP1 binding via one hydrogen bond (THR99) and four hydrophobic interactions (ALA95, LYS102, LEU47, LYS103) (also see-Supplementary **Fig. S3**). **(C-D)** Highlights the survival and death curve of *An. culicifacies*. To confirm the *in silico* predicted analysis we performed a standard WHO bottle assay to expose susceptible strains to pyrethroid (Deltamethrin) and recorded their survival/death in a time-dependent manner, and noticed a maximum survival of the exposed mosquitoes for up to 30 minutes; (E-G) Relative expression analysis of SAP1 in different tissues to deltamethrin exposure in the surviving adult female mosquito *An. culicifacies*. Statistical significance **p < 0.004; and ***p < 0.0004. ****p<0.00001 was calculated using student’s t-test and Mann-Whitney U test (n=represents the number of mosquitoes pooled for sample collection; N= number of replicates; C= Control; E= Exposed to Insecticide; Olf= Olfactory).

### AcSAP1 mediates body sensitization against insecticide exposure

Finally, to validate the expression of functional protein *AcSAP1* in the mosquito body (tissues), first, we cloned, expressed, and purified recombinant protein (**Fig4**: A, B; and Supplementary Text; **Fig. S4**) to generate anti-*AcSAP1* polyclonal antibodies. Immunolocalization studies using a polyclonal antiserum were conducted to examine the distribution of the *AcSAP1* protein throughout the body of the adult female mosquito *An. culicifacies*. Immunofluorescence and confocal microscopy revealed that *AcSAP1* is predominantly expressed in the sensilla of peripheral tissues. In control (un-exposed) females, *AcSAP1* expression was restricted to specific peripheral sensilla across the body. However, upon insecticide exposure, unexpectedly there was a remarkable increase in *AcSAP1* expression, extending to all sensilla of the peripheral tissues, especially surrounding the eyes (Fig.4C, D). These data suggested that the chemical nature and properties i.e. bad smell and toxicity of the insecticide may significantly impact distinct body tissues. We hypothesize a gradual interaction of insecticide with peripheral tissue viz. legs, wings, olfactory, and body sensilla may trigger a red alert signal taken forward to the eyes, and then possibly to the brain for necessary decision.

**Figure 4:**
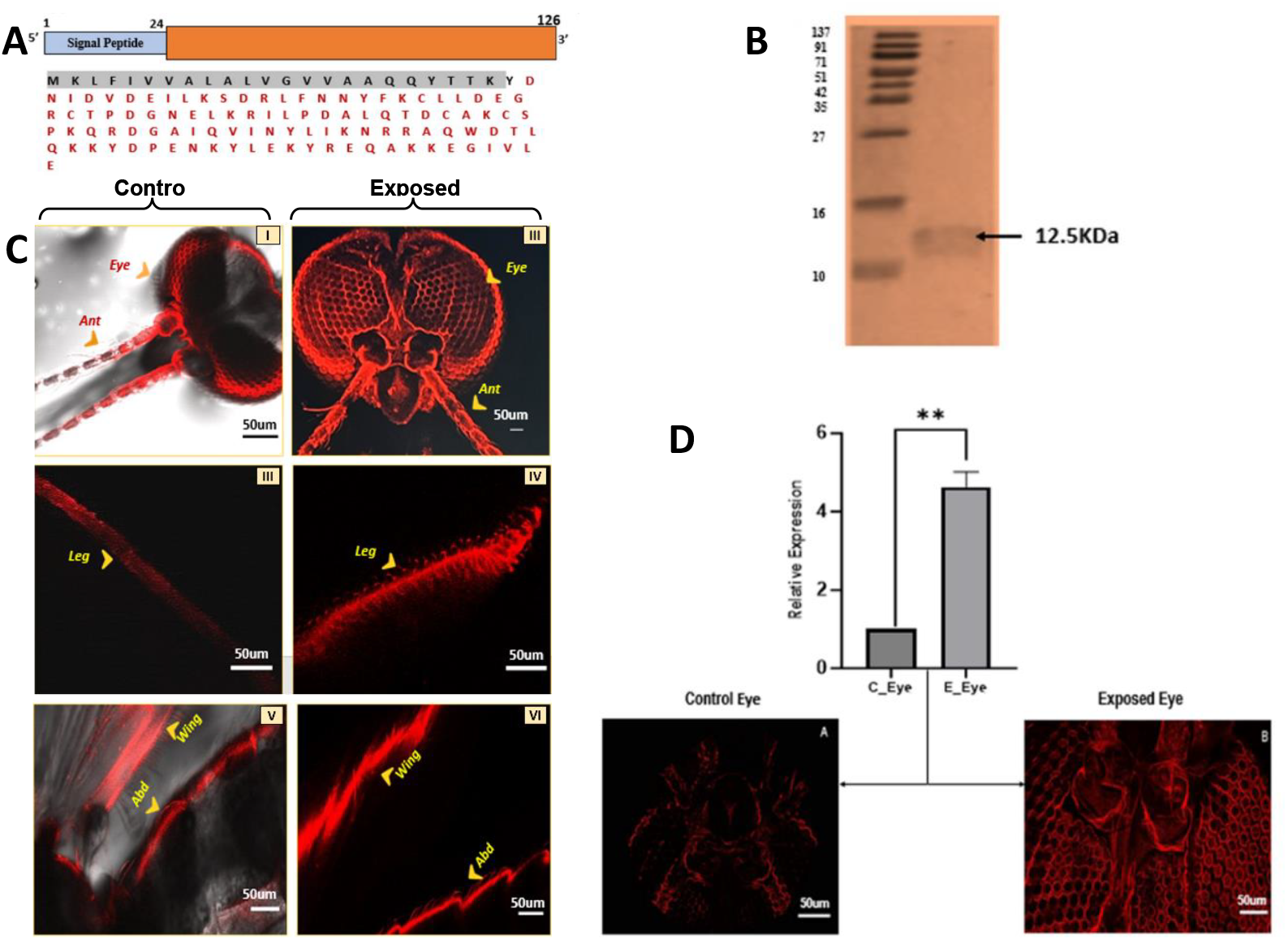
Confocal microscopy assay: **(A)** Schematic overview of *AcSAP1* primary structure: Coding region with single exon shown in orange and the signal peptide in blue. The size of each region is indicated in base pairs (bp). The deduced amino acid sequence of *AcSAP1* protein showing the predicted signal peptide (Black shaded), and the amino acids comprising OS-D in Red letters; **(B)** Western blot analysis of *AcSAP1* recombinant protein: Immunostaining was performed with a polyclonal antiserum against *AcSAP1*. Markers (M) size ranges from the top: 137, 91, 71, 51, 42, 35, 27, 16, 10kDa; **(C)** *AcSAP1* immunolocalization and expression in the peripheral tissues of the females *An. culicifacies*. Comparative assessment of cellular expression of *AcSAP1* in the control (Left images - I, III, V) and insecticide exposed (Right images -II, IV, VI) mosquito tissues. Quantitative change in the florescent intensity among two groups was also compared and evaluated for our confocal assays (see Supplementary **Fig. S5**); **(D)** Relative expression analysis of *AcSAP1* in eyes to deltamethrin exposure in the adult female mosquito *An. culicifacies*. Statistical significance **p<0.005 was calculated using student’s t-test and Mann-Whitney U test (n=represents the number of mosquitoes pooled for sample collection; N= number of replicates, Ant=Antennae/ Olfactory part, Abd= Abdomen sensilla);

## Discussion

While navigating towards the host, distinct phases of host-seeking activities are coordinated by concerted actions of different sensory organs (23). We hypothesize that a successful navigation trajectory may result in the integrated action of multiple, overlapping sequential events in peripheral sensory tissues i.e. (i) Olfaction-mediated detection of host over a distance *via* environmental cues (24), (ii) Gradual activation of legs and wings while approaching towards landing phase, and (iii) Eye engagement to finding/locating a suitable site to punch and suck the blood (25).

In the past decade, we and others have identified a group of chemosensory proteins (CSPs/SAPs) dominantly expressing in the peripheral sensory appendage i.e., olfactory and legs of the malarial vectors (3, 6). In *An. culicifacies*, the late-night upregulation of SAP transcripts suggested its role in modulating host-seeking properties (3) though the underlying mechanism remains unexplored. We tested whether the identified sensory appendage proteins (SAPs) play any role in host-seeking-associated activities. A reduced host attraction was more evident in the *AcSAP1* mRNA-depleted mosquitoes compared to the control mosquito groups, confirming SAP’s role in modulating host-seeking behavior.

Since insecticides are well known to play a vital role in the disruption of the navigation trajectory (26), we further tested the impact of insecticide exposure on SAP expression in different sensory organs of the laboratory-reared susceptible strain of *An. culicifacies* (27). A 3D model of the structure showed efficient binding of synthetic pyrethroid i.e. deltamethrin suggesting that the crucial interaction of SAPs could be key to the modulating mosquito’s behavioral responses. (28, 29). Survival for up to 30 minutes of exposure was found enough to evaluate the molecular response of SAP in the major peripheral tissues of the surviving mosquitoes (30, 31). Like other toxic chemical agents, insecticides also have distinct chemical properties *viz*. bad smell, toxicity, and irritancy, that harm our health (32-33), we hypothesized that SAPs may likely play a key role in the behavioral modification, that allows *An. culicifacies* to avoid the toxic environment (34). Compared to the control group, an elevated expression in the olfactory, legs, and wings, strongly supports the idea that SAPs possess the ability to recognize the intensity of the bad smell and/or toxicity. This ability could direct a pre-alert to sensitize the peripherally active sensory body system/organs

An elevated signal of SAP/antibody binding in all the peripheral tissues viz. olfactory system, legs, and wings as well as body parts evidenced insecticide-mediated body sensitization. However, surprisingly, a remarkable induction of the SAP expression in the eyes of the pyrethroid-exposed mosquitoes is an unusual observation that has not been previously reported in any mosquito species (35). These findings suggest that integrated coordination of SAPs could be central for guiding and managing the navigation trajectory (36), and if interfered with any toxic agent like insecticides, SAP may also ensure an early sensing alertness to detect and avoid insecticide response.

Summarily, based on our findings we propose that (i) a circadian-dependent modulation of SAP expression is crucial for managing the host-seeking associated navigation trajectory, and (ii) the ability to bind and recognize multiple properties of insecticides such as bad smell, toxicity, and irritant nature, could serve as a key first-line mechanism to avoid toxic response by integrating sensory information from different peripheral appendages. We hypothesize a deep understanding of SAP interaction with distinct chemical cues, and the mechanism of pre-alert strategies could be beneficial to designing a novel vector control tool **(Fig.5)**.

**Figure 5:**
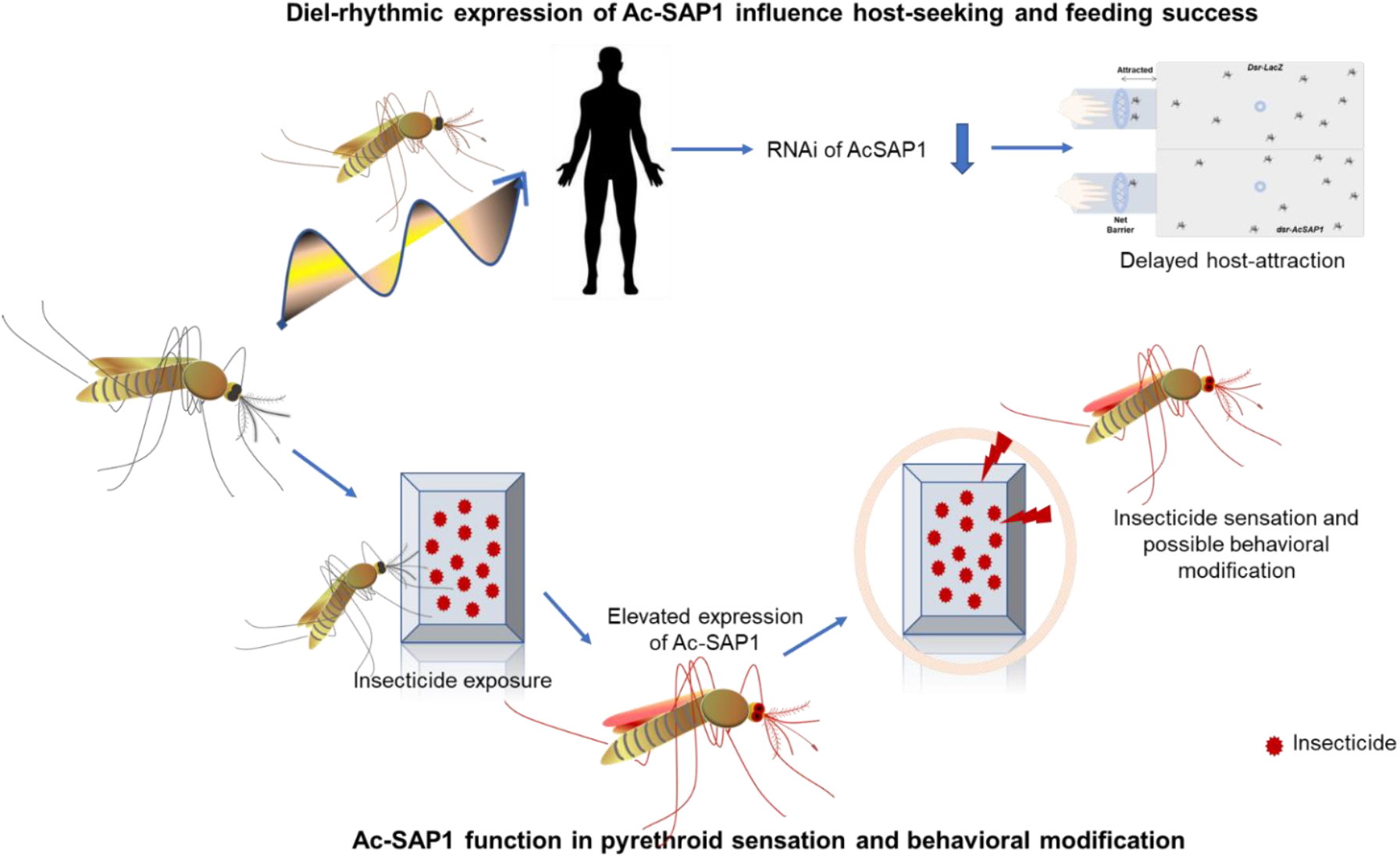
A proposed hypothesis of *AcSAP’s* role in host-seeking and behavioral modification properties: Host-seeking in *An. culicifacies* involve coordinated actions of sensory organs, including olfaction for detecting hosts, leg and wing activation for navigation, and eye engagement for locating blood-feeding sites. Sensory appendage proteins (SAPs) are crucial in this process, as evidenced by reduced host attraction in SAP-silenced mosquitoes. SAPs also play a role in insecticide response, showing elevated expression in peripheral tissues and eyes upon pyrethroid exposure, indicating their role in detecting and avoiding toxic environments. These findings suggest that SAPs integrate sensory information to manage navigation and insecticide avoidance, highlighting their potential as targets for innovative vector control strategies.

## Supporting information

Supplemental data sheet

## Acknowledgment

We extend our gratitude to the insectary staff for their rigorous efforts in mosquito rearing. We also appreciate Kunwarjeet Singh, and Lipun Pradhan for their technical support in the laboratory. We are thankful to GeNext Genomics Pvt Ltd., Nagpur, India for cloning and expression and generation of antibodies. We are grateful to Dr. Neetu Singh - Technician B, AIRF, JNU, New Delhi, India for confocal microscopy. Finally, we are highly thankful to ICMR-National Institute of Malaria Research, Delhi for extending the infrastructural and other logistic support for the project work.

## Authors contribution statement

Vaishali S., T.D.D., P.R., Vartika S., P.Y., T.S., G.T., G.S., and N.S.: Contributed to design and performing the experiments, data acquisition, writing and editing; Vaishali S., T.D.D., S.T., and R.D.: Conceptualization, and designed the experiments, Data analysis and interpretation, data presentation, contributed reagents/materials/analysis tools, wrote, reviewed, edited, and finalized M.S. All authors have read and agreed to the published version of the manuscript.

## Ethical Clearance

Necessary ethical clearance was taken for relevant research work.

