## Supplemental data sheet for "Sensory Appendage Protein triggers alarm to pyrethroid in Indian malarial vector Anopheles culicifacies"

**Supplementary Table-1:** Homology-based identified seven putative CSPs from olfactory RNAseq data of adult female mosquito *An. culicifacies*

| S.N. | RNAseq Transcript-ID | Feeding Status | Label | AC-ID | Ag Homology | Ag-ID |
|---|---|---|---|---|---|---|
| 1. | Transcript_1808 | Naive | CSP-1 | ACUA010232 | Chemosensory Protein (E-value: 6.00E-51) (Identity: 96.70%) | AGAP008055 (CSP-3) |
| 2. | Transcript_447 | Naive | CSP-3 | ACUA004458 | Chemosensory Protein (E-value: 7.00E-71) (Identity: 96.70%) | AGAP008059 (CSP-1) |
| 3. | Transcript_702 | Naive | SAP-3 | ACUA017151 (SAP-3) | Chemosensory Protein (E-value: 1.00E-29) (Identity: 90.90%) | AGAP008054 (SAP-3) |
| 4. | Transcript_12 | Naive | SAP-1 | ACUA018714 (SAP-1) | Chemosensory Protein (E-value: 2.00E-40) (Identity: 99.73%) | AGAP008051 (SAP-1) |
| 5. | Transcript_1446 | 30m_PBF | CSP-5 | ACUA013563 | Chemosensory Protein (E-value: 5.00E-80) (Identity: 99.40%) | AGAP008058 (CSP-5) |
| 6. | Transcript_6551 | Naïve | SAP-2 | ACUA013409 | Chemosensory Protein (E-value: 2e-76) (Identity: 98.18%) | AGAP00852 (SAP-2) |
| 7. | Transcript_2682 | 30hr PBM | CSP-6 | ACUA008309 | Chemosensory Protein (E-value: 1.00E-66) (Identity: 99.9%) | AGAP001303 (CSP-6) |

PBF= Post blood-feeding;

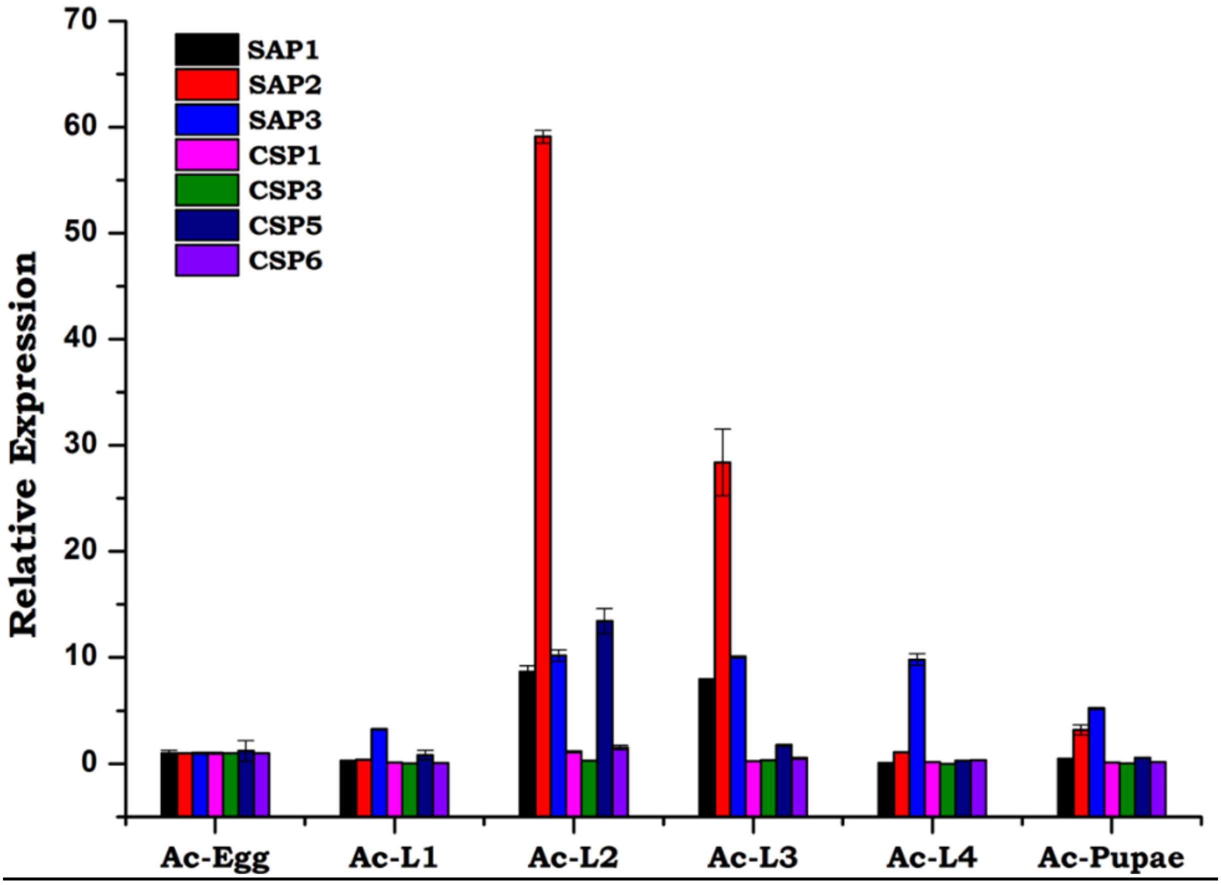


**Figure S1: Developmental expression profiling of seven CSP/SAP members in the mosquito *An. culicifacies:*** Relative gene expression during aquatic development, showing relative expression of identified members of CSPs/SAP transcripts in the mosquito *An. culicifacies*. The egg was considered the control for all test samples. L1-4 (larval stages) (n=10, N=3).

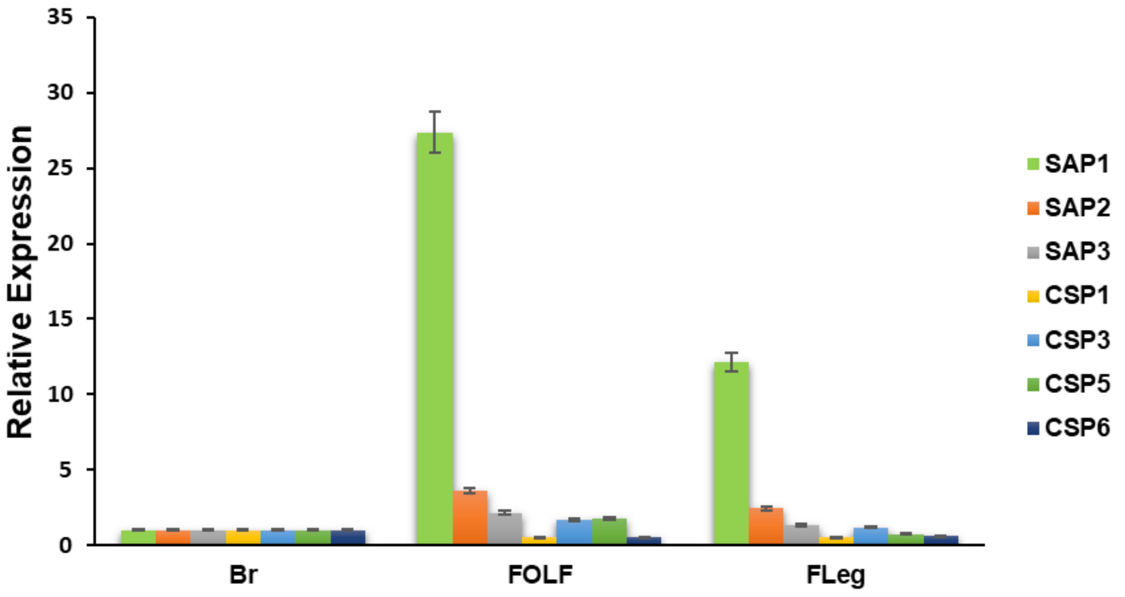


**Figure S2**: Tissue-specific Expression analysis: CSP/SAPs differentially expressed in peripheral sensory organs: Olfactory (OLF) and Legs were dissected from 3-4 days old naïve adult female mosquitoes *An. culicifacies*. Graphs show abundant expression of SAPs in the olfactory and leg tissues. Data were compared with female brains as a control (n = 25, N=3).

**<u>Supplemental Table 2:</u>** *In-silico* prediction and comparative analysis of the physio-chemical nature of SAP proteins

| S. No. | Property | SAP1 | SAP2 | SAP3 |
|---|---|---|---|---|
| 1 | Accession no. (VectorBase) | ACUA018714 | ACUA013409 | ACUA017151 |
| 2 | Sequence length | 126 | 127 | 126 |
| 3 | Molecular weight | 14617.84 | 14764.96 | 14563.70 |
| 4 | Theoretical pI | 8.29 | 5.34 | 8.57 |
| 5 | Instability index | 36.06 | 34.69 | 29.67 |
| 6 | Aliphatic index | 93.65 | 84.57 | 84.37 |
| 7 | Grand average of hydropathicity (GRAVY) | -0.550 | -0.598 | -0.685 |
| 8 | Extinction coefficients (units of M-1 cm-1, at 280 nm measured in water) | 16180 | 16180 | 17670 |
| 9 | Subcellular localization | Cytoplasmic | Cytoplasmic | Cytoplasmic |

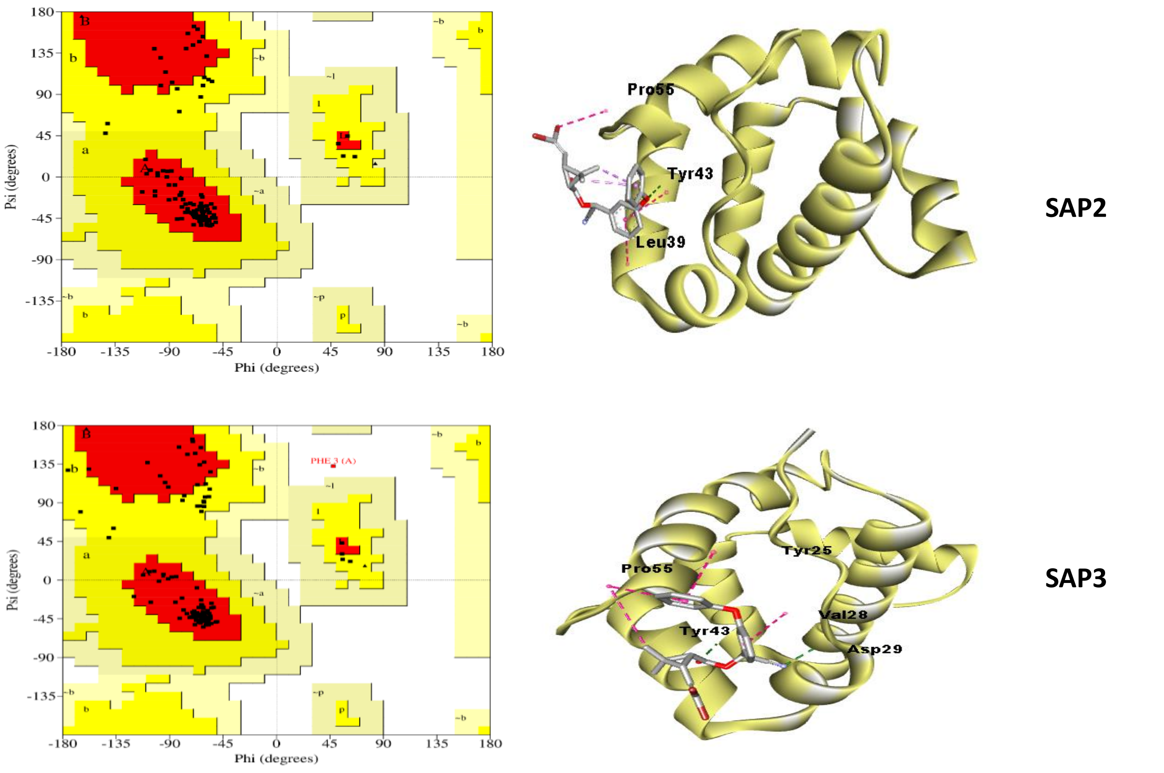


**<u>Figure S3: *In-silico* 3-D structural modelling, and molecular docking analysis of SAP/insecticide interaction:</u>** Ramachandran plots for SAP2 and SAP3 proteins**.** In this figure red color indicates the core region, yellow indicates allowed regions, lemon indicates generously allowed and white is disallowed region. Docked complexes of SAP proteins with Deltamethrin: SAP2 is forming 1 hydrogen bond (TYR25) and 3 hydrophobic interactions (LEU39, PRO55) while SAP3 participating in two hydrogen bonds (ASP29, TYR43) and three hydrophobic interactions (TYR25, VAL28, PRO55). Following is the color of bond representation: Green- Hydrogen Bond, Pink- Hydrophobic interactions.

**Extended supplementary Methodology Text**

**Cloning, bacterial expression, and protein purification:** The first few amino acids were observed as signal peptides, and removed before designing cloning primers (Supplemental Fig4A). SAP1 protein was cloned into the bacterial expression vector pET28a (+) (Novagen, Madison, WI) between the Nco1-Xho1 restriction sites. The SAP1 recombinant plasmid was then extracted, verified by PCR (FigS4B)/sequencing, and then transformed into E. coli BL21competent cells. A single colony was grown overnight in 50 ml LB broth (including 50 µg/ml kanamycin). Five liters of LB medium were inoculated with the 50 ml overnight culture at 37°C for 2–3 h until the absorbance at OD 600 reached 0.6. The protein expression was then induced for 22 hours using IPTG with a final concentration of 1 mM at 18°C. The bacterial cells were harvested by centrifugation (16,000 rpm, 30 min), resuspended in a lysis buffer (buffer A - 50 mM Tris-HCl Ph 8.0, 300 mM NaCl, 10% glycerol), lysed by sonication for 20 min with 2 min interval and centrifuged again (16,000 rpm, 30 min). The supernatant was collected and mixed with equilibrated (buffer A) Ni-NTA beads and kept on a rotator for O/N. Next day O/N Ni-NTA beads packed in column and washed with 20 mM Imidazole in a buffer. Finally, the protein was eluted with 300 mM Imidazole. Collected elution fractions were pooled, concentrated up to 5-6 ml and loaded on Hi-Load 16/600 Superdex 200 column AKTA purification. Collected fractions analyzed by 15% SDS-PAGE (Fig S4C-D).

**ELISA**: Recombinant SAP1 (200 ng) was coated using 1X PBST buffer in 96 well microtiter plates at 4 °C overnight. Unbound protein was washed with 1X PBST and blocking was done with 2% BSA. Rabbit raised polyclonal primary antibody of different titer (1.5ng, 3.15ng, 6.25ng, 12.5ng, 25ng, 50ng, 100ng and 200ng) was incubated for 1 h at 37 °C with the coated protein. Unbound antibody was washed and incubated with HRP conjugated anti rabbit Alexa 594 secondary antibody (1:10,000). After washing, TMB detection reagent (Himedia, India) was added and the reaction was stopped using stop buffer (0.3N $NH_2SO4$). Absorbance was detected at 450 nm in microplate reader (Varioskan, Termo, USA).

**Western blot analysis**: Purified SAP1 was separated on 15% SDS-PAGE and then transferred to a PolyVinylidene Fluoride (PVDF), membrane using a machine (Genscript ALLIANZ bio). The membrane was blocked with 2% BSA in 1X TBST at room temperature and washed three times with PBST (5 min each). The blocked membrane was then incubated with the purified rabbit anti-SAP antiserum (2ug/ml) for 1 h at room temperature and washed with TBST. Subsequently, the membrane was incubated with an anti-rabbit secondary antibody (1:10K). After repeated washing, the membrane was developed with TMB and visualized with Enhanced Chemiluminescence detection reagents (GE Healthcare).

**A**

**Signal Peptide**

MKLFIVVALALVGVVAAQQYTTKYDNIDVDEILKSDRLFNNYFKCLLDEGRCTPDGNELKRILPDALQTDCAK
CSPKQRDGAIQVINYLIKNRRAQWDTLQKKYDPENKYLEKYREQAKKEGIVLE

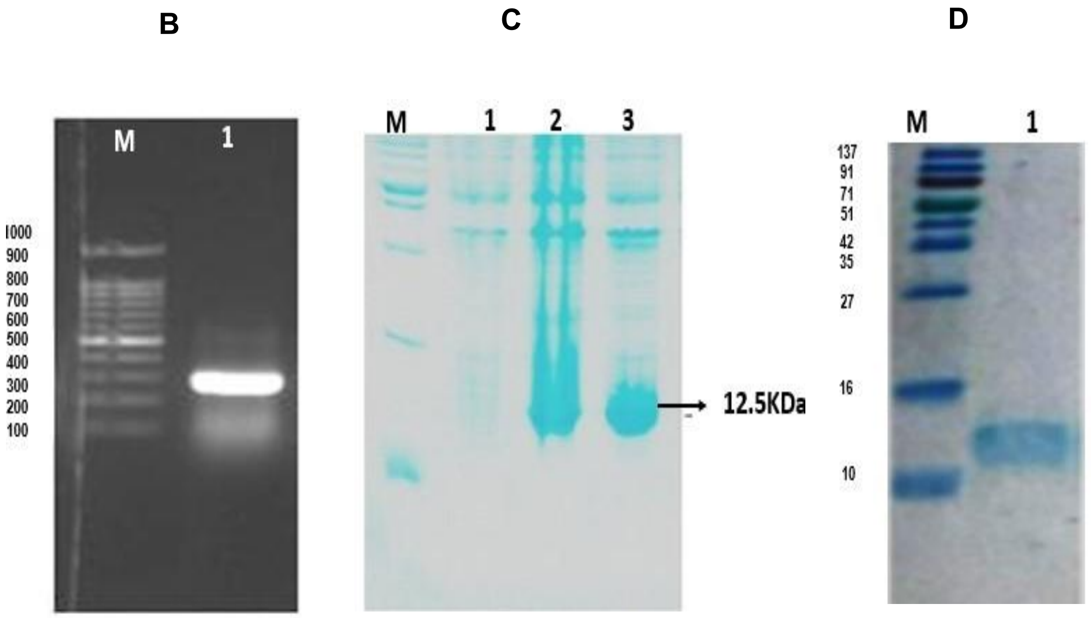


**Figure S4:**

(A)Deduced amino acid sequence of *AcSAP1* protein showing the predicted signal peptide (Black), the pro-peptide (pink), and the amino acids comprising OS-D in Red letters. (B) PCR and clone verification. (C) Protein induction (M -Marker, 1- uninduced, 2- whole cells, 3- Supernatant after sonication. (D) Ni-NTA purified protein was gel filtered by size exclusion chromatography HiLoad 16/600 Superdex 200 column

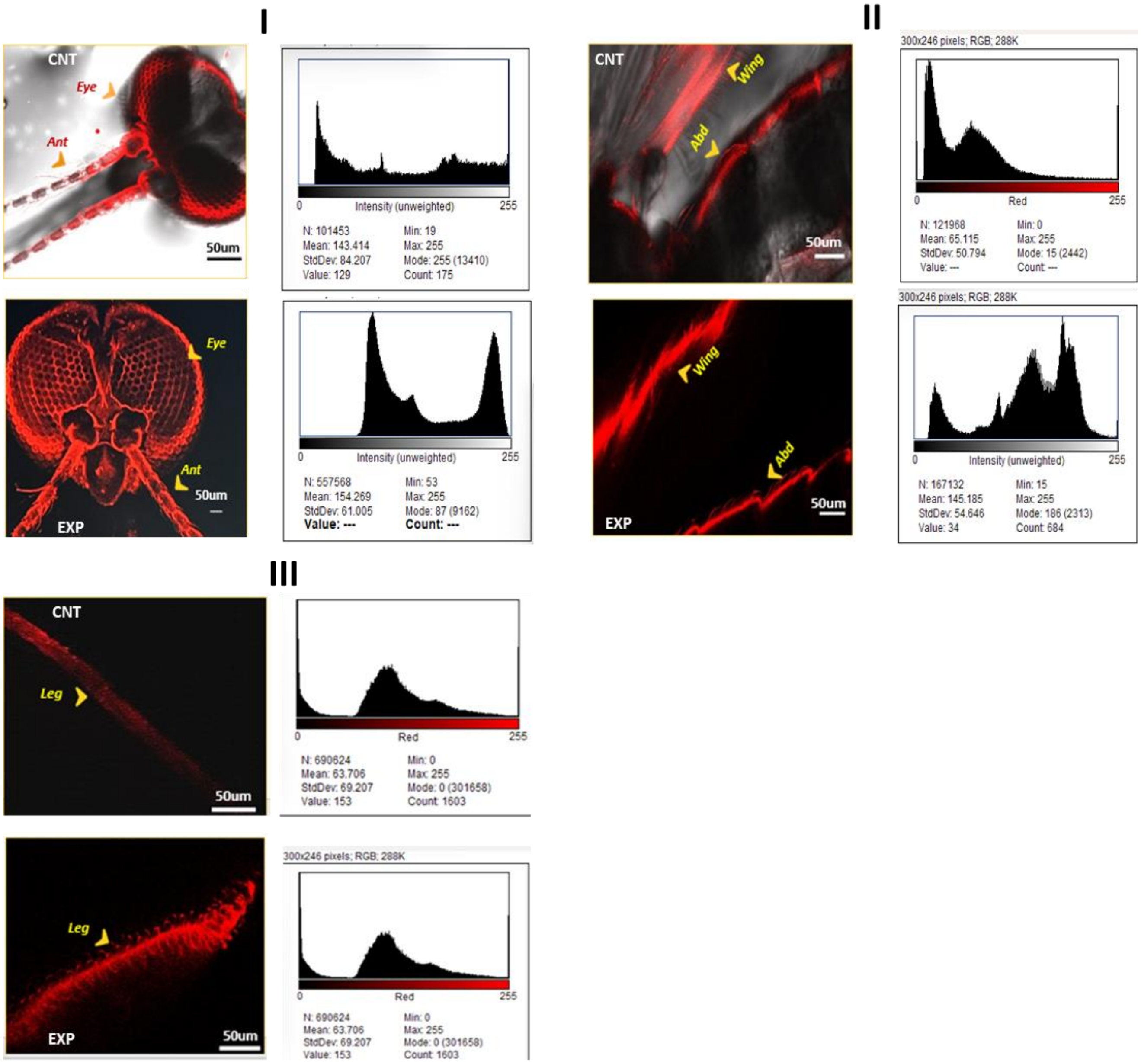


**Figure S5: Confocal images with Intensity Histogram variation:** Anti-SAP1 antibody labelled images were captured under a confocal microscope, and corresponding histograms were generated to compare the fluorescent intensity variation among Control (CNT) and insecticide-exposed (EXP) mosquito's sensory tissues viz. Panel-I [Antennae (Ant) and Eyes (Eye)]; Panel-II [Wing and Abdomen (Abd)]; Panel-III [Legs]
